# Stochastic population models to identify optimal and cost-effective harvest strategies for feral pig eradication

**DOI:** 10.1101/2023.03.08.531659

**Authors:** Peter W. Hamnett, Frédérik Saltré, Brad Page, Myall Tarran, Matt Korcz, Kate Fielder, Lindell Andrews, Corey J. A. Bradshaw

## Abstract

Eradicating feral pigs from island ecosystems can assist in restoring damaged biodiversity values and protect commercial industries such as agriculture. Although many feral pig eradications have been attempted, management decisions are often led by practitioner experience rather than empirical evidence. Few interventions have been guided by population models to identify harvest rates necessary to achieve eradication within a specified time frame, nor have they applied data on control effort and cost to evaluate the relative cost-effectiveness of proposed control strategies. We used effort and cost data from a feral pig-control program on Kangaroo Island, South Australia over 17 months to derive functional-response relationships between control effort (hours pig^-1^) and pig abundance for four control methods: (*i*) ground-based shooting, (*ii*) trapping with remote triggers, (*iii*) poison baiting, and (*iv*) thermal-assisted aerial culling. We developed a stochastic Leslie matrix with compensatory density feedback on survival and fertility to project population trajectories from an initial population (*N*_0_) of 250 female pigs with an estimated island-wide carrying capacity (*K*) of 2500 over 3 and 10 years for populations subjected to an annual harvest of 35% to 95%. We built functional-response models to calculate annual effort and cost for six cull scenarios across all harvest rates. We derived total cost and effort over 3- and 10-year projections from the sum of annual cost and effort within the projection intervals. Pig populations were reduced to < 10% *N*_0_ based on harvest rates > 70% and 50% for culls of 3- and 10-year duration, respectively. In all scenarios except ‘trapping only’, the total cost to reduce population to ≤ 10% of *N*_0_ decreased with increasing harvest proportion, with lower total costs incurred over 3 years compared to 10 years. The simulations suggest that the most cost-effective approach for most scenarios is to maximise annual harvest and complete eradication effort over the shortest periods.

## Introduction

Feral pigs (*Sus scrofa*) — a species derived from both wild boar and domesticated pig breeds — are major pests in much of their introduced range. Feral pigs cause a wide range of environmental, economic, and social damages, including the destruction of native vegetation and productive land, competition with/predation on native fauna and livestock, spread of other invasive species and pathogens (Barrios-Garcia and Ballari 2012, Bengsen et al. 2014), and emissions of organic carbon from soil disturbance (O’Bryan et al. 2022). Within their native range of Eurasia and northern Africa (IUCN 2022), the species spans a vast area encompassing diverse climatic and environmental conditions that, along with their omnivorous diet and high reproductive rate, make them a successful invasive species (Lewis et al. 2017). The species has been introduced widely outside of its native range through accidental release and abandonment of domestic herds, and intentional release to establish populations for bushmeat and sport hunting (Courchamp et al. 2003, Long 2003). Feral pigs now occur on all continents (except Antarctica) and many oceanic islands, with future range expansion predicted in North America, South America, Africa, and Australia (Lewis et al. 2017). In Australia, feral pigs occupy about 40% of the mainland and offshore islands (Lapidge et al. 2012), with a total population estimated at 13.5 million (95% confidence interval from 3.5 to 23.5 million) (Hone 1990a, Choquenot 1996).

Globally, pigs threaten at least 672 taxa, including 638 taxa in their non-native range (Risch et al. 2021). In Australia, feral pigs are a key threatening process identified under the *Environment Protection and Biodiversity Conservation Act 1999*, with impacts on at least 148 nationally threatened species and 8 threatened ecological communities (Commonwealth of Australia 2017). The number of threatened species impacted by feral pigs in South Australia has not been quantified, but impacts to biodiversity and primary production are recognised through a pest-animal declaration under the *Landscape South Australia Act 2019*, placing legal obligation on landholders to destroy feral pigs (Government of South Australia 2022).

Despite the breadth of environmental degradation by feral pigs, the associated costs are not readily quantified and economic estimates of their impact typically focus on productivity losses and management costs (Cowled and Lapidge 2004, McLeod 2004). In the USA, losses and damages from feral pigs are conservatively estimated at US$2.95 billion over the period 1983 to 2017 (Diagne et al. 2021). In Australia, costs since 1960 are estimated to range between US$9.54 billion considering all available reported costs, and US$0.73 billion when only highly reliable costs are considered (Bradshaw et al. 2021). Mean costs for management and economic losses for all invasive alien species worldwide are conservatively estimated at US$26.8 billion year^-1^ for the period 1970 to 2017, with costs roughly doubling every 6 years over this period and projected to continue rising as the expanding footprint of global transport, trade, and development create new opportunities for invasions (Diagne et al. 2021).

For the biodiverse and relatively intact native vegetation on Kangaroo Island in South Australia (Australia’s third-largest island at ∼ 443,000 ha), feral pigs represent an ongoing hazard to many threatened and endemic species (Masters et al. 2011). A sow and boar were first introduced to Kangaroo Island in January 1803 by French explorer Captain Nicolas Baudin (1754–1803) at Anse des Sources, which early settlers later renamed Hog Bay (Fig. 1) after the pig population became established there (Cooper 1954). Since their initial introduction, the feral pig population has grown through both natural recruitment and supplementation by intentional and accidental releases (Masters et al. 2011). Pigs had disappeared from Hog Bay and Antechamber Bay by 1880–1890, but occurred in the Wilson River District and west of Parndana, including Starvation Creek, South West River, and Flinders Chase by the 1950s (Pullar 1953, Cooper 1954). Between 1984 to 2019 the population was estimated to be stable, with about 5400 individuals distributed over about 1350 km^2^ on the western end of the island, despite sporadic efforts to reduce their numbers (Masters et al. 2011, Southgate and Florance 2018). Today, feral pigs are estimated to cost the local economy ∼ AU$1 million year^-1^ (Primary Industries and Regions South Australia 2020).

**Figure 1.**
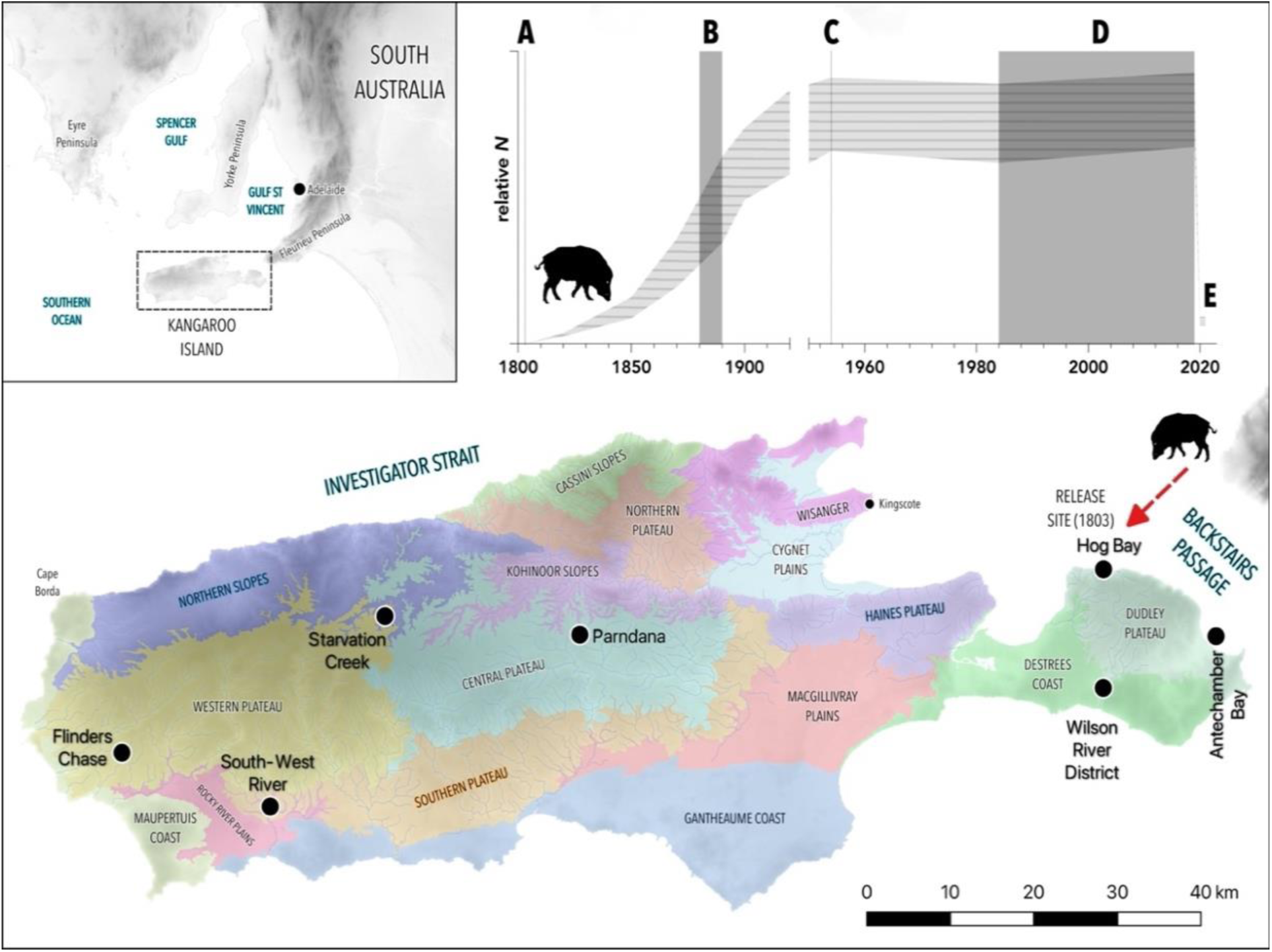
Kangaroo Island, showing its major biophysical land zones, and sites of historical importance in the spread of feral pigs. The inset graph shows an approximate reconstruction of relative population size across the island (*N*) inferred from historical reports: (A) release at Hog Bay, (B) expansion to south and east on Dudley Peninsula, (C) present at western end of island but no longer in the east, (D) formalisation of monitoring indicates population is stable, (E) bushfires reduce population.

The catastrophic bushfires of summer 2019–2020 (Boer et al. 2020) reduced the pig population to about 500 individuals, presenting a rare opportunity to achieve eradication. However, their high reproductive rate — average litter size ≥ 8, physiologically capable of producing 2 litters in 14 months, and often reaching sexual maturity within the first year of life (Oliver et al. 1993) — allows pig populations to recover quickly following density reduction or stochastic events (Choquenot 1996), so immediate action was required. The Australian and South Australian Governments allocated ∼ AU$2.66 million over three years to achieve eradication by winter 2023 (Primary Industries and Regions South Australia 2020). At the time of publication, the estimated pig population was < 30 individuals following the commencement of the eradication program (Primary Industries and Regions South Australia 2021).

Where possible, eradication is preferrable to continued management of invasive species because it curtails ongoing costs associated with lost productivity and management (Bomford and O’Brien 1995, Baxter et al. 2008). However, eradication is only feasible when certain criteria can be met, including: (*i*) rate of pest removal must exceed the rate of recruitment at all densities; (*ii*) no immigration or ability to remove immigrants quickly; and (*iii*) all individuals in the population must be susceptible to the available control methods. Additionally, desirable criteria include: (*iv*) the ability to detect the pest at low densities; (*v*) cost of eradication is justified compared to ongoing population control; and (*vi*) sufficient socio-political and economic support to achieve success (Bomford and O’Brien 1995). Given available resources, methods, and the extent of feral pig occurrence in Australia, continent-wide eradication is not considered possible, although eradication of isolated island populations might be (Commonwealth of Australia 2017), as shown by successful pig eradications around the world that have predominantly occurred on small islands (e.g., Miller and Mullette 1985, Statham and Middleton 1987, Kessler 2002, Parkes and Murphy 2003, Cruz et al. 2005, Parkes et al. 2010). It has yet to be demonstrated if all criteria for successful eradication of feral pigs on Kangaroo Island can be satisfied, but the recent eradications of fallow deer (*Dama dama*) and feral goats (*Capra hircus*) (Masters et al. 2018) on the island inspire confidence that pig eradication is also achievable.

A variety of lethal (e.g., trapping, snaring, ground- and aerial-shooting, poisoning, dogs) and non-lethal methods (e.g., fertility control, fencing, repellents, diversionary feeding, translocation, and disruption) are used to control feral pigs, although the feasibility and efficacy of each depend on local conditions, legal constraints, social acceptability, and management objectives (Campbell and Long 2009, Massei et al. 2011, Bengsen et al. 2014). On Kangaroo Island, private landholders have historically used shooting, trapping, and dogs to control pigs, while government agencies have trialled alternative methods such as poisoning with sodium fluoroacetate (1080/PIGOUT®), as well as investigating the use of radio-collared ‘judas’ pigs to improve pig detection of remaining herds (Masters et al. 2011). Methods currently used on Kangaroo Island include ground-based shooting, poisoning with sodium nitrite (HOGGONE®), trapping, and thermal-assisted aerial culling — a novel approach to aerial culling that uses thermal imagery to improve detection (Cox et al. 2022, Bradshaw et al. 2023).

Despite increasing global costs from damages and management of invasive species, reduction and eradication campaigns are often *ad hoc*, informed by personal experience, anecdotal evidence, and best guesses by land managers rather than knowledge derived from research, monitoring, or formal assessment (Pullin et al. 2004, Sutherland et al. 2004). While this can lead to success in some instances (Parkes et al. 2010, Holmes et al. 2015), such an approach is susceptible to bias and might lead to suboptimal implementation, wasted resources, and even failure (Cook et al. 2010, McMahon et al. 2010). Long-term and repeated failure to achieve project objectives can undermine support for spending on invasive-species management, particularly where control methods have low social acceptability (e.g., aerial 1080 baiting or using hunting dogs) (Massei et al. 2011, Sinclair et al. 2019). Optimisation of invasive species strategies to maximise cost-effectiveness and probability of success are there for important for managing the growing costs of invasive species management.

Simulation models to project population responses to different management regimes can be used to assess the effectiveness of different approaches against project parameters (e.g., timeframe, cost, population threshold) (Baker and Bode 2021). While the use of models to guide eradication is not a new concept, the application of population modelling to feral pig management on Kangaroo Island has not been reported. Similarly, the application of operational effort data to estimate locally relevant functional responses and predict future management costs at later stages of control programs is novel. Functional responses describe the relationships between *per capita* efficiency of a predator’s ability to catch and consume prey as a function of prey density (Holling 1959), and they are suitable for describing changes in efficiency of methods to control pest animals (Hone 1990b). For a particular control method, the form of the functional response can be inferred by observing changes in effort (e.g., hours pig^-1^, pigs hour^-1^) relative to some measure of pig abundance or density (e.g., pigs km^-2^, proportion of total population remaining), and is influenced by the effect of declining density on rate of offtake, whether there is a threshold density above which *per capita* rate of offtake becomes saturated, and if there are refuges within the landscape in which animals can escape detection (Choquenot et al. 1999). The availability of operational records from feral pig control on Kangaroo Island allows locally relevant functional response to be estimated.

Our aim was to design a set of management scenarios that will lead to the eradication of feral pigs in Kangaroo Island. More specifically, we (*i*) built a stochastic population model, (*ii)* projected the reduction of the pig population based on different methods of control, and (*iii*) estimated the relative costs of employing the different methods available. We used operational cost and effort observations from pig control on Kangaroo Island to estimate functional responses for four control methods (ground shooting, trapping, poison baiting, and thermal-assisted aerial culling). We applied these functional responses to estimate the total effort and cost required to achieve population reduction to ≤ 0.1 of the initial population (*N*_0_ = 250) for six control scenarios, including four scenarios relying on one method only, and two scenarios that applied all four control methods in fixed and varying proportions to simulate generic best practice. Although the target timeframe for pig eradication on Kangaroo Island is 3 years, we applied each control scenario over timeframes ranging from one to ten years to identify if efficiency gains could be achieved by applying controls at lower intensity and over a longer duration. We applied all harvest scenarios at annual harvest proportions ranging from 0.35 to 0.95. We hypothesise that a reduction in the population to ≤ 0.1*N*_0_ is achievable using available control methods within the 3-year timeframe and AU$2.66 million budget. Ultimately, we address the following questions: What will it cost to achieve eradication using a particular control regime? Which regime will achieve success for the lowest cost or effort?

## Methods

### Study site

Kangaroo Island is Australia’s third-largest island, with an area of approximately 443,000 ha (Fig. 1) (Robinson et al. 1999, Tyler et al. 2002). Large tracts of native forest and shrubland remain, particularly in the west and south of the island, at the coastal fringe, and on roadsides, accounting for approximately 53% of total landcover (Willoughby et al. 2018). Approximately 68% of remaining natural vegetation is protected within Conservation Reserves and Wilderness Protection Areas, accounting for around 32% of the island’s area (Robinson et al. 1999). Dryland agriculture and plantation forestry account for 35% and 3% of landcover, respectively, and are major components of the Kangaroo Island economy (O’Neil et al. 2017, Willoughby et al. 2018).

### Stochastic population model

We constructed a female-only, post-breeding Leslie (age-structured) matrix model in R (R Core Team 2022) to project annual population growth (Caswell 2001). The model assumes a sex ratio of 1:1 (Snow et al. 2019) and equal probability of survival between males and females. In Australia, few pigs are suspected to live beyond 5 years (Choquenot 1996), although longevity up to 12–14 years has been reported (Snow et al. 2019). We set the maximum age at 6 years, although probability of survival in age class *n*_6_ > 0 (see Table 1) allowing for a few individuals to live beyond 6 years.

**Table 1.** Mean fertility and survival values and their standard deviation (SD) for all age classes used in the stochastic model.

| vital rate | mean | SD |
| --- | --- | --- |
| *fertility* (daughters) |  |  |
| juvenile ($f_1$) | 0.963 | 0.150 |
| yearling ($f_2$) | 2.338 | 0.178 |
| adult ($f_{3-5}$) | 2.925 | 0.590 |
| senescent ($f_6$) | 2.835 | 0.590 |
| *survival* |  |  |
| juvenile ($s_1$) | 0.300 | 0.068 |
| yearling ($s_2$) | 0.400 | 0.073 |
| adult ($s_{3-5}$) | 0.660 | 0.033 |
| senescent ($s_6$) | 0.580 | 0.033 |

Given a maximum age of 6 years, the deterministic matrix **A** is:

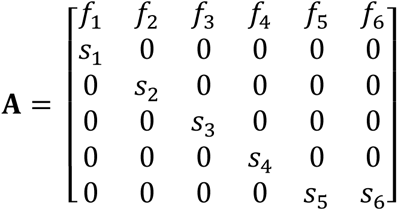

where *f_x_* = age specific fertility (the number of female offspring per individual per year in age class *x*) and *s_x_* = age specific survival (the probability of surviving from age *t* to *t*+1). For an initial population *N*:

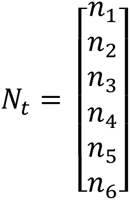

where *n_x_* = number of females in age class *x* at time *t*, population growth can be projected as:

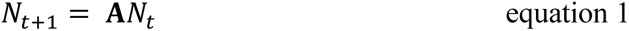

The mean and standard deviation of age-specific vital rates (survival and fertility) are required to project stochastic population growth. These are unreported for the Kangaroo Island pig population and data are insufficient to calculate these from harvest records (e.g., Skalski et al. 2005). In lieu of available local estimates, we used compiled mean pig vital rates for poor, intermediate, and good years based on food availability and winter climate from European studies, mostly in Germany and Eastern Europe (Bieber and Ruf 2005) (Appendix S1: Table S1). We selected intermediate-year values as means for age-specific survival and fertility in age classes *n*_1_ to *n*_5_, assuming these would most closely represent conditions on Kangaroo Island. We assigned the senescent age class (≥ *n*_6_) survival and fertility values for poor years. We calculated standard deviations for survival and fertility using values from poor, intermediate, and good years (Table 1).

At each time-step and for each age class, we accounted for stochastic variation in the model by randomly resampling from Gaussian and *β* distributions around the means of fertility and survival probabilities, respectively (Table 1).

### Population projections

To produce a vector of females per age class (*N*_0_), we calculated the population’s stable stage distribution from the deterministic matrix **A** (Caswell 2001) and multiplied this by the initial population size of 250 (females only) (B. Page, Adelaide, South Australia, personal observation). We used *N*_0_ as the initial population in all simulations. We assumed that the carrying capacity (*K*) was 5000 individuals. Because the Leslie matrix projects changes in the number of females only, we adopted *K* = 2500 for the female-only model. Systematic estimates of the Kangaroo Island pig population have not been reported but, prior to the 2019–2020 bushfires, the population was estimated at 675 and 5400 individuals based on densities of 0.5 to 4 pigs km^-2^ observed elsewhere in Australia (Choquenot 1996, Masters et al. 2011). Monitoring over the period 2009 to 2018 estimated a stable population (Masters et al. 2011, Southgate 2013, Southgate and Florance 2018, Primary Industries and Regions South Australia 2020) and discussions with local wildlife managers on Kangaroo Island supported the estimated *K* and pre-bushfire population of *∼* 5000.

At each time increment of the projection, both survival and fertility probabilities in all age classes were recalculated using a modifier to simulate compensatory density feedback. We modified survival and fertility (*S*_mod_ and *F*_mod_, respectively) as:

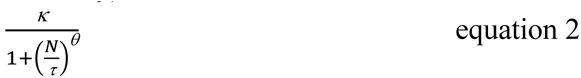

where *N* = population size, *κ*, *τ*, and *θ* are constants (*κ* = 1,*τ* = 2500, *θ* = 3) such that applied survival and fertility probabilities decrease as the population approaches *K*. We defined these constants arbitrarily through iterative changes to the modifiers until the population projection returned the expected response (Fig. 2). Initially, we applied density feedback only to survival, but the influence of high fertility resulted in population projections exhibiting exponential growth at the upper confidence interval (Appendix S2: Fig. S1). We resolved this biological issue by applying the modifier to both survival and fertility, assuming that compensatory density-feedback on fertility is representative of both variation in litter size with declining maternal condition (Frauendorf et al. 2016) and reduced neonatal survival as resource competition increases as the population approaches *K*.

**Figure 2.**
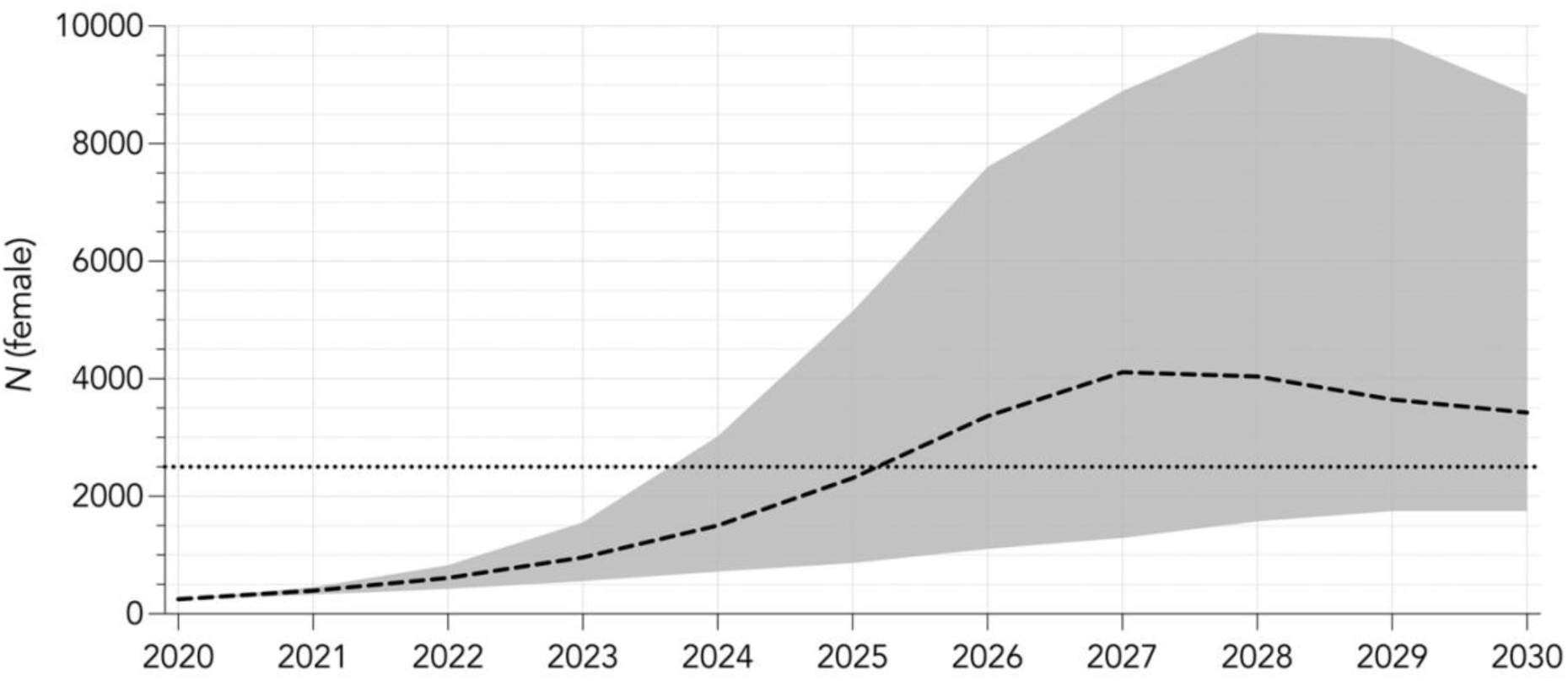
Stochastic population growth in an untreated population from an initial population (*N*_0_) of 250 females over 10-year projection interval with compensatory density feedback applied to both survival and fertility. Black dashed line indicates median population growth from 1000 iterations, grey shaded area indicates 95% confidence interval, and black dotted horizontal line indicates estimated carrying capacity (*K* = 2500 females).

We first simulated a ‘control’ stochastic projection (1000 iterations) to examine median projected growth in an untreated population, i.e., a population not subjected to any density-reduction measures. We projected population growth from *N*_0_ over 10 years, incorporating annual stochasticity and density feedback on survival and fertility as described above. We then built a constant proportional cull model to examine the influence of annual harvest rate on the proportion of population remaining over two different projection intervals — 3 and 10 years — and identify threshold harvest rates for achieving reductions to *N*_0_ ≤ 0.1 (≤ 25 females; 1% of *K*). The threshold is based on the lower end of the ‘50/500’ rule where < 50 *N*_e_ (effective population size) is considered prone to inbreeding depression and extinction (Frankham et al. 2014). We selected the 3-year projection interval to align with the Government of South Australia’s target timeframe for the eradication of feral pigs from Kangaroo Island (Primary Industries and Regions South Australia 2020). We also selected a 10-year projection interval arbitrarily (but conceivably within a management-relevant timeframe) for comparison. For each year of the projection interval, the constant proportional cull model reduced the population by the same proportion, from 0.2 to 0.95 in increments of 0.05. For each harvest proportion, we repeated the model over 1000 iterations, incorporating stochastic variation in fertility and survival as described above. For all iterations and harvest rates, we recorded the minimum projected proportional population size after each time increment. From these, we calculated median minimum proportional population size and 95% confidence intervals for all harvest rates at both projection intervals.

Because the 2019–2020 bushfires killed most of the feral pigs on Kagaroo Island, we repeated 3- and 10-year annual cull simulations on a founding population *N*_0_ = 2500 to examine differences in minimum constant proportional cull rates to achieve population reduction to *N*_0_ ≤ 0.01 (≤ 25 females; 1% of *K*) and total median costs under the more likely circumstances that land managers start density reduction on a pig population at or close to carrying capacity. We provide the results of this additional analysis as Supplementary Material (Appendix S4).

### Pig-control data

The Government of South Australia Department of Primary Industries and Regions provided pig-control data from 29 April 2020 to 7 December 2021. The data included outcomes from pig control completed by a range of control actors, including government employees (Primary Industries and Regions, South Australia, Kangaroo Island Landscapes Board and National Parks and Wildlife Service, South Australia), pest-control contractors, non-government organisations, and members of the public. Each record included the date, time, geographic coordinates, number of pigs killed, and the control method used. Most records included the amount of effort (hours) expended per event and name of the individual or organisation who did the control. Additional details were only reported for some pigs, including the age and sex of individuals killed and/or number of individuals evading control, and so we could not include these in the model.

### Pig-control methods

The data included four main pig-control methods: (*i*) shooting, (*ii*) thermal-assisted aerial culling, (*iii*) trapping, and (*iv*) poison baiting. Additional control methods/causes of death reported included vehicle strike, dogs and ‘other’. Ground shooting was done through active searching (e.g., tracking, spotlighting, thermal imagery), opportunistic encounters, and shooting at sites where feed had been deployed as an attractant. Shooting records did not specify the approach taken, but we inferred this from knowledge of the preferred method of individual pig controllers through discussion with project staff (M. Korcz, Kingscote, South Australia, personal observation) and estimated that 58.8% of total shooting effort was through shooting with free feeding.

Thermal-assisted aerial culling data were collected during two culls done from 18–30 March 2021 and 20 August 2021–24 September 2021. HeliSurveys (helisurveys.com.au) was engaged to provide the helicopter, pilot, and thermal camera operator, while the marksman was a Government of South Australia employee. The helicopter travelled at a height of 50–100 m above ground and at between 15–25 knots ground speed while searching.

Trapping was done using the MINE (manually initiated nuisance elimination) Trapping System (Jager Pro 2022), a remotely triggered, corral-style trap (diameter ∼ 10.5 m, height ∼ 1.7 m). The traps are connected to the mobile communication network and use motion-sensing cameras to detect movement within the trap. Operators are notified by text or email when motion is detected, real-time images are reviewed, and a decision made either to trigger the trap or wait. To minimise neophobic avoidance behaviour, trap deployment and activation occurred in a staged manner over several days, including delivery of trap components to the site, gradual assembly of the corral, and deployment of grain (attractant).

Poison baiting was done using HOGGONE® meSN® microencapsulated sodium nitrite bait (Animal Control Technologies Australia 2020). Baiting involves a staged process, like that used for trapping, whereby free feeding encourages pigs to congregate at the bait site over several days before placebo baits are introduced to train feeding from the bait dispenser and ultimate deployment of toxic bait.

### Efficiency

To estimate changes in efficiency of control methods relative to proportion of pigs remaining, we calculated specific relationships between the proportion of pigs remaining after each control event and efficiency for individual control methods. Efficiency (hours pig^-1^) of the *i*^th^ event (*E_i_*) is:

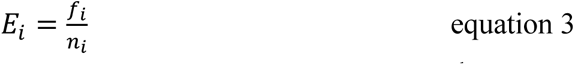

where *n_i_* = number of pigs killed in the *i*^th^ event and *f_i_ =* effort (hours) expended in the *i*^th^ event. For thermal-assisted aerial culling, multiple kill events were often recorded per flight (e.g., several distinct groups of pigs encountered during the flight), but the effort was only recorded as total flight time per flight. As such, we counted each flight as a unique event and calculated efficiency as the total number of pigs killed flight^-1^ divided by the flight duration. We could not calculate event-specific efficiency for events where effort was not reported, nor could we calculate it for events where cause of death was reported as anything other than by using one of the four main control methods.

Because we do not know the true number of pigs in the total population, we calculated change in proportion of pigs remaining relative to the total number of pigs killed during the study period (29 April 2020 to 7 December 2021), plus the estimated number remaining on Kangaroo Island (*n* = 200) at the end of the study period (Primary Industries and Regions South Australia 2021). Therefore, the proportion of the feral pig population remaining (*N_p_*_,*i*_) after the *i*^th^ event was:

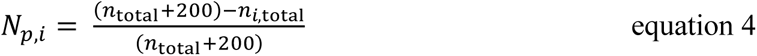

where *n*_total_ = total number of pigs killed during the collection period (*n* = 757) and *n_i_*_,total_ = total number of pigs killed up to and including the *i*^th^ event. We included all reported pig kills in the total number killed, including events where effort was unreported or cause of death was reported as anything other than one of the four main control methods, to allow calculation of proportional population change over time.

### Functional responses

To compare model performance, we fitted logarithmic, exponential, and linear models to the relationship between proportion of pig population remaining and effort pig^-1^ for each control method. We assumed that the efficiency of all control methods decreases as population density declines, following a Type II functional response (Hone 1990b, Choquenot et al. 1999) resulting from decreasing probability of ‘capture’ (detection/destruction) (Caley and Ottley 1995).

Contrary to this assumption, raw data for trapping, shooting, and poisoning exhibited trends of increasing efficiency with decreasing proportion of pigs remaining. For each control method, we therefore identified outliers with the assistance of the contributing individual participant or organisation. We removed all such outlying data contributed by the individual or organisation and refit the models. This resulted in satisfactory relationships in the expected direction for trapping and poisoning, but not for opportunistic shooting or shooting with free feeding. We therefore aggregated the data for opportunistic shooting and shooting with free feeding, resulting in a relationship following the expected direction.

We used Akaike’s information criterion (*w*AIC) and the information-theoretic evidence ratio to compare model performance to identify which model (logarithmic, exponential, linear) best fit the abundance-efficiency functional response. Differences between model performances were negligible across all control methods (Table 2). We chose the exponential model to apply in the subsequent stages of effort and cost estimation (see below) because it conforms best to the expected Type II functional response.

**Table 2.** Method-specific maximum efficiency (hours pig^-1^) derived from operational observations.

| **control technique** | **minimum effort kill<sup>-1</sup>**<br>**(hours $\text{pig}^{-1}$)** |
| --- | --- |
| thermal-assisted aerial culling | 0.128 |
| shooting | 0.222 |
| trapping | 0.333 |
| poisoning | 0.308 |

### Costs

Generic costs that were applicable to more than one control method included: (*i*) *labour* — AU$36.87 person^-1^ hour^-1^, based on South Australia Public Sector Award OPS4 classification (Commissioner for Public Sector Employment 2017). (*ii*) *ammunition —* AU$4.00 pig^-1^ based on AU$2.00 bullet^-1^ at a rate of 2 bullets pig^-1^ allowing for misses and sight zeroing. (*iii*) *feed grain* — AU$14.00 day^-1^ site^-1^ based on deployment of 10 kg grain at AU$1.4 kg^-1^ and assuming grain-deployment effort of 1 hour day^-1^. We assumed the same effort rate for deployment of placebo bait and toxic bait. We excluded several generic costs from the cost calculations, including general administrative overheads (office space and equipment, project administration, office-based staff, community/stakeholder engagement etc.), vehicle lease and fuel, and costs associated with deploying and maintaining trail cameras for monitoring pig activity. We did not include cost of firearms in the method-specific costs assuming firearms can be employed across multiple control methods, e.g., destruction of pigs caught in traps, ground-based and aerial shooting. Furthermore, the cost of firearms is negligible compared to costs for labour and other materials or equipment.

We summarised method-specific costs as:

(i) *Thermal-assisted aerial cull* — project initialisation cost AU$12,146, including crew mobilisation (pilot and camera operator) and helicopter from Jindabyne, New South Wales, project data management, and initial fuel delivery. Crew and helicopter flight time, including wages, fuel, and maintenance, cost AU$2482.66 day^-1^, with average flight effort of 3.7 hours day^-1^. Additional daily costs include helicopter crew meals and accommodation, and government marksman labour, meals, and accommodation. Helicopter crew meals and accommodation cost AU$420 day^-1^ crew^-1^. Government marksman labour cost assumed the marksman was engaged in cull-related activities on a full-time basis for the duration of the cull. Labour cost was accrued at the OPS4 hourly labour rate in increments of 7.5 hours day^-1,^ based on 37.5 hours week^-1^, equating to AU$276.53 day^-1^. Marksman accommodation and meals were charged separately to those of the helicopter crew, costing AU$125 day^-1^. A fuel-resupply charge AU$1246 was incurred every 30 days, equivalent to 111 hours of flight effort based on average flight effort of 3.7 hours day^-1^.
(ii) *Shooting* — costs largely comprised hourly labour and bullets per pig, as described in the generic costs above. Additionally, free-feeding cost was accrued at AU$8.23 hour^-1^ (AU$14 hour^-1^ multiplied by 0.588) based on the proportion of shooting effort estimated to have occurred with free feeding.
(iii) *Trapping* — we assumed effort per trapping event to be constant, estimated at 29.875 hours trap^-1^ based on mean observed effort of 9.875 hours event^-1^ plus estimated additional 20 hours event^-1^ trap set-up time that was not included in reported effort (M. Korcz, Kingscote, South Australia, personal observation). We calculated grain cost at AU$138.25 event^-1^ based on mean observed effort multiplied by generic free-feeding cost hour^-1^. Labour cost AU$1101.49 event^-1^ based on 29.875 hours trap^-1^ multiplied by the generic labour cost hour^-1^. Jager Smart Traps cost AU$9500 each (M. Tarran, Adelaide, South Australia, personal observation). A single trap has capacity to complete an average of 10 trapping events year^-1^ based on estimated deployment time of 4–6 weeks event^-1^ (M. Korcz, Kingscote, South Australia, personal observation), equating to annual effort of 298.75 hours trap^-1^ year^-1^ (29.875 hours trap^-1^ × 10 events year^-1^). We assumed traps were reusable for the duration of the projection intervals such that additional traps were only purchased in subsequent years if the model projected an increase in trap numbers over time, with the number purchased in any year equal to the number required minus the sum of traps purchased in all previous years.
(iv) *Baiting* — we assumed the mean observed baiting effort of 12.692 hours event^-1^ to be constant for cost estimation. Labour cost was AU$467.97 event^-1^ based on mean observed baiting effort multiplied by the generic labour cost hour^-1^. Cost of grain for free-feeding was AU$107.69 event^-1^ (mean observed baiting effort minus 5 hours event^-1^ multiplied by the generic free-feeding cost hour^-1^). We deducted 5 hours because grain is replaced by placebo baits (4 hours event^-1^) and toxic baits (1 hour event^-1^) in the final stages of baiting. Bait dispensers cost AU$485 each and have capacity for 6 placebo bait or toxic baits. Multiple dispensers were used at baiting sites in some instances, with a mean rate of 1.45 dispensers site^-1^ during the collection period. An individual dispenser or set of dispensers were capable of servicing 28.76 events year^-1^, assuming constant deployment at 365 days year^-1^, 1 hour effort day^-1^, and 12.692 hours event^-1^. As with trapping, we assumed that bait dispensers were reusable for the duration of the projection intervals and additional dispensers only purchased in subsequent years to make up shortfall if dispensers purchased in previous years did not satisfy the number of traps required. Placebo baits cost AU$224 dispenser^-1^ event^-1^ (AU$14 each at 6 day^-1^ dispenser^-1^ for 4 days). Toxic baits cost AU$156.00 dispenser^-1^ event^-1^ (AU$26 each at 6 day^-1^ dispenser^-1^ for 1 day).

### Culling scenarios

We estimated effort and cost for six different culling scenarios, comprising four scenarios in which 100% of the harvest quota was achieved by each of the four control methods (i.e., shooting, thermal-assisted aerial culling, trapping, and poison baiting) individually. The remaining two scenarios applied the four control methods in combination, simulating generic integrated pest-management approaches. These included an equal proportion harvest scenario where each method removes 25% of the annual harvest quota, and a relative cost-proportional-allocation scenario that used all four control methods in varying proportions weighted in favour of the most cost effective control method, recalculated at each time interval according to the following equation:

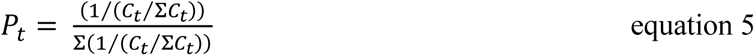

where *P_t_* is a 1 × 4 matrix with elements 1 to 4 being the proportion of annual offtake assigned to thermal-assisted aerial culling, shooting, trapping, and poisoning at time *t,* and *C_t_* is a 1 × 4 matrix with elements 1 to 4 being the total cost of thermal-assisted aerial culling, shooting, trapping, and poisoning at time *t*, if each method were used to complete 100% of the required offtake.

### Cost and effort estimates

We applied the six different culling scenarios at varying harvest proportions over projection intervals of 3 and 10 years for 1000 iterations, as per the constant proportional cull simulations described above. At each time increment for all simulations, we projected population growth and calculated the total number of pigs to be harvested by multiplying the projected population by the harvest proportion. We then subtracted the total number of pigs to be harvested from the total population at that time increment to give the population remaining after harvest, which we then multiplied by the stable stage distribution to give the number of pigs per age class for population projection in the next time increment.

For all scenarios, we determined the efficiency of harvest at each time increment by applying efficiency rates (equation 3) relative to the proportion of pigs remaining prior to harvest. Due to the asymptotic shape of the exponential functional response models, minimum effort per kill (hours pig ^-1^) for all control methods was limited according to the maximum observed efficiency calculated from operational records (Table 2), such that unrealisitically high efficiencies derived from the functional response model were not applied.

Proportion of pigs remaining (P) was:

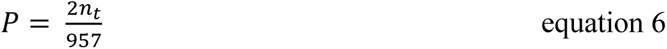

where *n_t_* is the number of females remaining at the beginning of the time increment, multiplied by 2 (to represent total population of male and females), divided by the population size used to determine the efficiency rates (*n* = 957). We calculated effort (hours) by multiplying the total number of pigs to be harvested by the relevant method’s proportion-specific efficiency rate.

At each time increment, we calculated the cost to reduce the population by the required annual harvest proportion by applying method-specific costs as defined above. For each iteration, we recorded cost and effort per year, as well as total cost and effort for the projection interval.

For each scenario, we used the outcomes after all iterations to calculate median and 95% confidence intervals of cost and effort per year, and totals for both projection intervals. For each cull scenario, we compared cost estimates generated using the most cost-effective harvest rates to the Government of South Australia operational budget for feral pig eradication to test if they could be achieved with the existing funding allocation.

## Results

### Untreated population projection

The deterministic matrix (without stochastic variation) produced an instantaneous rate of exponential change (*r*) of 0.344 (*λ* = *e^r^* = 1.411) and mean generation length of 2.9 years. Incorporating stochastic variation, the median projected population overshot carrying capacity (i.e., > 2500 females, Fig. 2) after 6 years, reaching maximum size of 16.4 times the initial population (*N*_6_ = 4111; 95% confidence intervals: 1474 and 9,865) after 7 years, before declining to 13.9 times the initial population (*n* = 3408; 95% confidence intervals: 1761 and 9,459) by the end of the 10-year projection interval (Fig. 2).

### Constant proportional cull

Constant annual harvest proportions of ≥ 0.7 and ≥ 0.5 were projected to achieve reduction to 0.1*N*_0_ (25 females) after 3 and 10 years, respectively (Fig. 3). Annual harvest proportions ≤ 0.35 produced negligible population change after either projection interval (< 1.2% and < 6% over 3 and 10 years, respectively).

**Figure 3.**
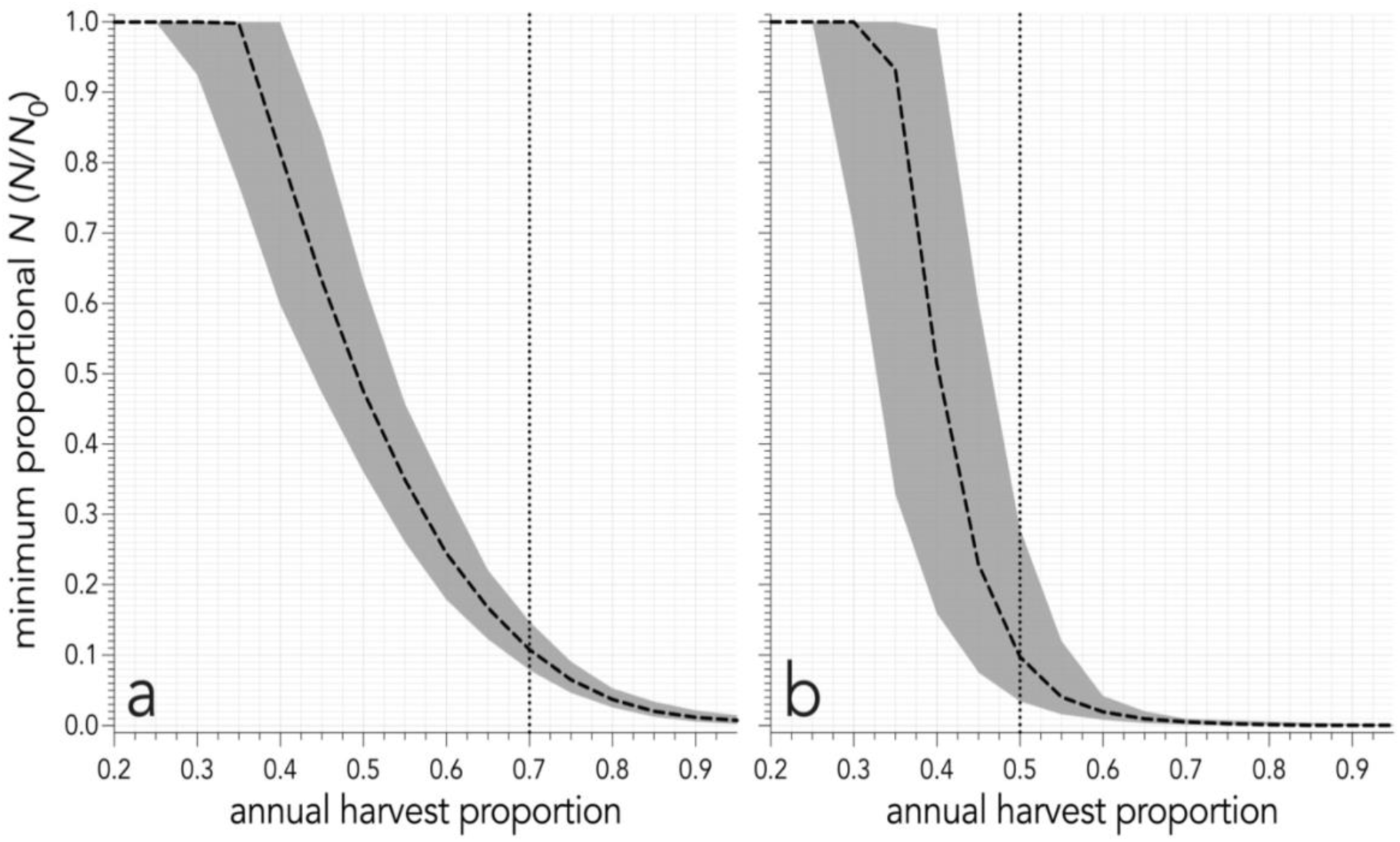
Proportion of initial population (*N =* 250) remaining after (a) 3 years and (b) 10 years of annual harvest at rates ranging from 0.2 to 0.95, increasing in increments of 0.05. Harvest targets all age classes equally. Black dashed line represents median proportion of population remaining along with its 95% confidence intervals (grey shaded area), vertical dotted lines represent harvest threshold required to reduce *N* to ≤ 0.1*N*_0_.

### Effort abundance relationships

The logarithmic model was top-ranked for trapping (Fig. 4), but differences between other models were negligible (Table 3). The exponential model was top-ranked for shooting, being 1.29 times (1/0.77) more likely to describe the relationship than the logarithmic model and 3.62 times more likely than the linear model. The linear model was top-ranked for both thermal-assisted aerial culling and poisoning, although differences between ranking of other models were negligible.

**Figure 4.**
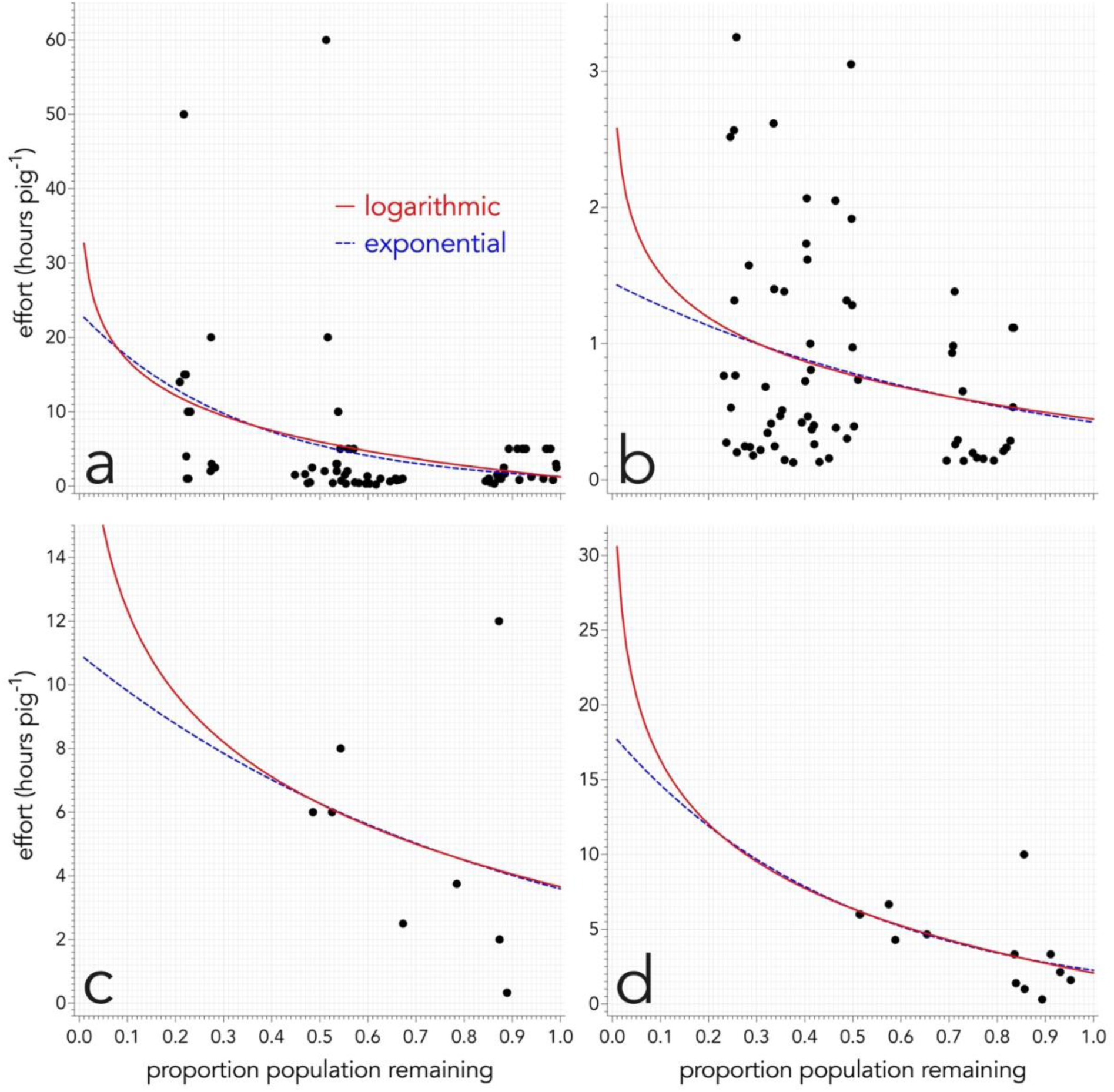
Effort per kill (hours pig^-1^) relative to proportion of pigs remaining for (a) shooting, (b) thermal-assisted aerial culling, (c) trapping, and (d) poisoning. Black dots represent individual events. Solid red and dashed blue lines represent the line of best fit for logarithmic and exponential models, respectively.

**Table 3.** Comparison of models (*log* = logarithmic, *exp* = exponential, *lin* = linear) fitted to the relationship between effort pig^-1^ and proportion of pig population remaining for trapping, shooting, thermal-assisted aerial culling, and poisoning. Bold *w*AIC indicate the top-ranked model for each control method.

| Control method | *w*AIC |  |  | evidence ratio |  |  |
| --- | --- | --- | --- | --- | --- | --- |
|  | *log* | *exp* | *lin* | *log:exp* | *exp:lin* | *log:lin* |
| trapping | 0.3361 | 0.3345 | **0.3294** | 1.00 | 1.02 | 1.02 |
| shooting | 0.3776 | 0.4877 | **0.1346** | 0.77 | 3.62 | 2.80 |
| thermal-assisted aerial culling | **0.3232** | 0.3292 | 0.3476 | 0.98 | 0.95 | 0.93 |
| poisoning | 0.3315 | **0.3214** | 0.3471 | 1.03 | 0.93 | 0.95 |

We derived parameters for estimating the required effort per pig killed (hours pig^-1^) from an exponential model of the form:

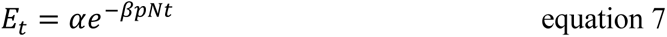

where *E_t_* is the efficiency (hours pig^-1^) at time *t*, *pN_t_* = proportion of pigs remaining, *e* is the exponential constant and *α* and *β* are constants unique to each control method (see Appendix S3, Table S1).

### Costs

Estimated cost to achieve the reduction target varied widely among control scenarios, annual harvest rates, and durations. Shooting with a 0.95 annual harvest rate was most cost-effective scenario over both 3- and 10-year projection intervals and cost AU$162,214 (AU$140,351 – AU$191,109) and AU$262,667 (AU$230,978 – AU$312,000), respectively. The least cost-effective scenarios to achieve successful reduction over 3 years (thermal assisted aerial culling; 0.7 annual harvest rate) and 10 years (thermal assisted aerial culling; 0.5 annual harvest rate) cost AU$428,634 (AU$370,097 – AU$492,383) and AU$896,918 (AU$669,194 – AU$1,253,317), or 264% and 341% more expensive than the most cost-effective combination of control scenario and annual harvest rate over the 3- and 10-year projection intervals.

The annual harvest rate of 0.95 produced the lowest total median costs for all individual control scenarios over both 3-year and 10-year projection intervals, except for trapping that had the lowest total median costs with a 0.85 harvest over both 3- and 10-year intervals (Fig. 5a,b).

**Figure 5.**
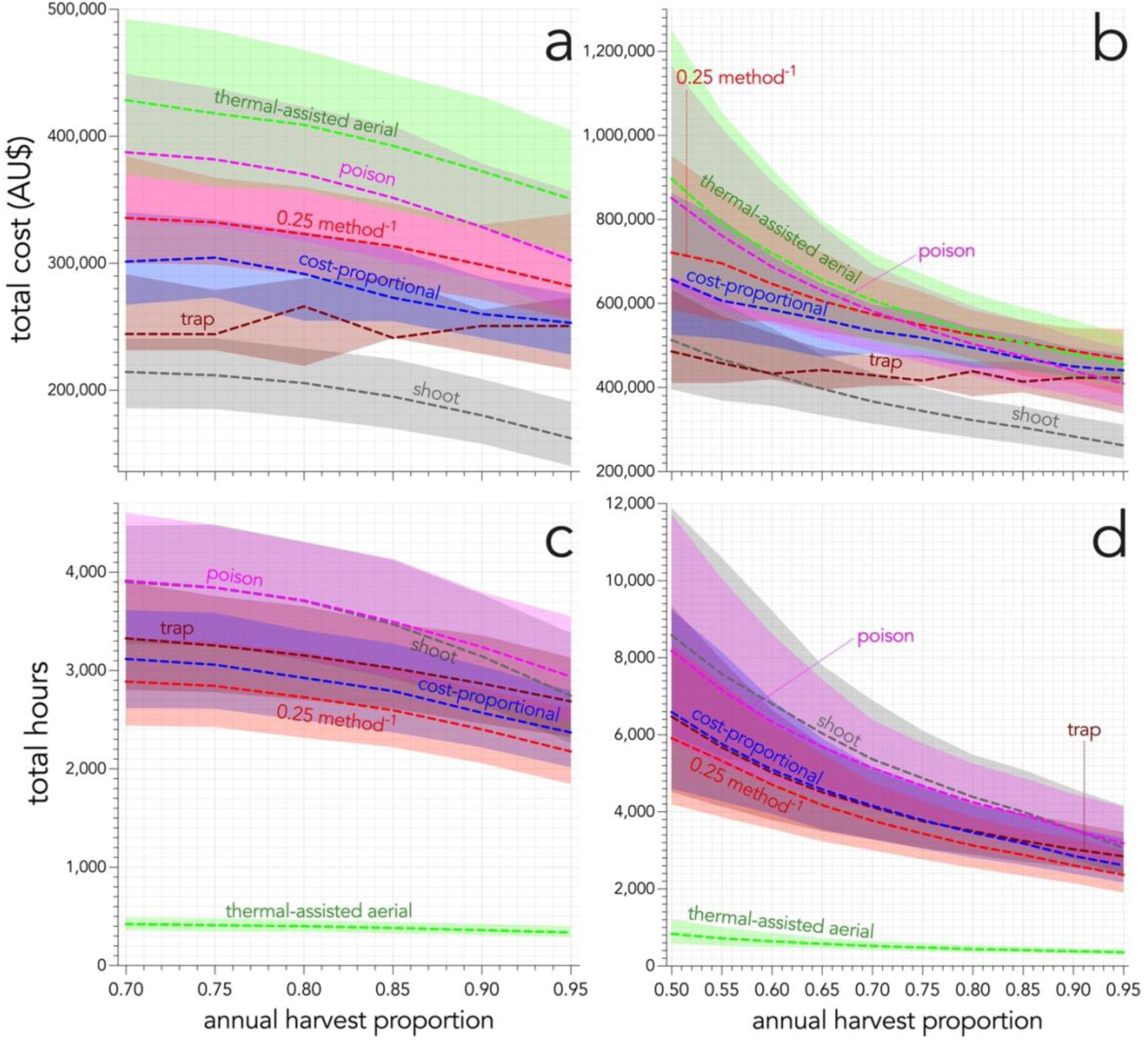
(a, b) Projected total cost (shaded areas = 95% confidence intervals) per cull scenario over (a) 3 and (b) 10 years, and (c, d) projected total effort (hours) per cull scenario over (c) 3 and (d) 10 years under increasing harvest proportions. Minimum harvest required to achieve reduction to *N* ≤ 0.1*N*_0_ is 0.7 (a, c) and 0.5 (b, d). **thermal-assisted aerial** = thermal-assisted aerial culling; **0.25 method^-1^** = 25% harvest per method; **cost-proportional** = cost-proportional allocation.

Additionally, both culling scenarios that simulated generic integrated pest management were most cost-effective at an annual harvest of 0.95. After 3- and 10-year projection intervals, the 25% harvest per method scenario produced total median costs of AU$282,173 (AU$256,277 – AU$339,075) and AU$468,649 (AU$383,614 – AU$539,355), respectively, or 174% and 178% more expensive than the most cost-effective scenario, whereas the relative cost-proportional-allocation scenario produced total median costs of AU$253,236 (AU$228,176 – AU$275,976) and AU$440,792 (AU$410,060 – AU$479,862), respectively, or 156% and 168% more expensive than the most cost-effective scenario.

At the maximum annual harvest (0.95), the reduction target (≤ 0.1*N*_0_) was achieved after just 1 year (∼ 0.08*N*_0_; 20 females remaining) and reduction to < 0.02*N*_0_ (< 5 females remaining) was achieved after 2 years. Comparison of costs estimated after 1 and 2 years with total median cost estimates after the full 3- and 10-year projection intervals therefore identified cost savings that could be achieved if control activities ceased after 1 or 2 years rather than continuing for the duration of the 3- or 10-year projection interval. Shooting remained the cheapest control method, with a median minimum cost of AU$99,397 (AU$90,001 – AU$107,854) after 1 year, and AU$141,682 (AU$124,692 – AU$162,497) after 2 years. Cessation of shooting at 0.95 harvest after 1 year saved AU$62,817 (39% reduction), and AU$163,270 (62% reduction) compared to total median costs accrued over the 3- and 10-year projection intervals. If shooting ceased after 2 years, the project saved AU$20,532 (13% reduction) and AU$120,985 (46% reduction) relative to the 3- and 10-year cost estimates. Similar cost savings occurred in all other culling scenarios.

Of the original ∼ AU$2.66 million budget for the feral pig eradication on Kangaroo Island over three years from July 2020 to June 2023, ∼ AU$1.8 million was allocated to operational costs (Primary Industries and Regions South Australia 2020), with remaining funds directed to program management and administration. Annual operational budget for each year of the program is AU$561,229 (2020–2021), AU$551,000 (2021–2022), and AU$686,000 (2022–2023). Comparing the operational budget to annual and cumulative cost estimates over the 3-year projection interval gave estimated total costs to achieve the reduction target (≤ 0.1*N*_0_) that were less than the actual amount allocated to operational costs in the first year of the program for all cull scenarios when applied at harvest rates > 0.7, the threshold for achieving the population target within 3 years (Fig. 5a). Additionally, for all control scenarios applied at 0.95 annual harvest, the estimated cost for continuing population reduction in the second and third year decreased substantially relative to estimated first-year costs. Second-year costs decreased between 57% (shooting) and 81% (thermal-assisted aerial culling), and third-year costs decreased between 79% (shooting) and 91% (thermal-assisted aerial culling) relative to first-year costs. By comparison, actual allocated funding decreased slightly (2%) in the second year but increased to 120% of the first-year funding allocation in the third year (Fig. 6).

**Figure 6.**
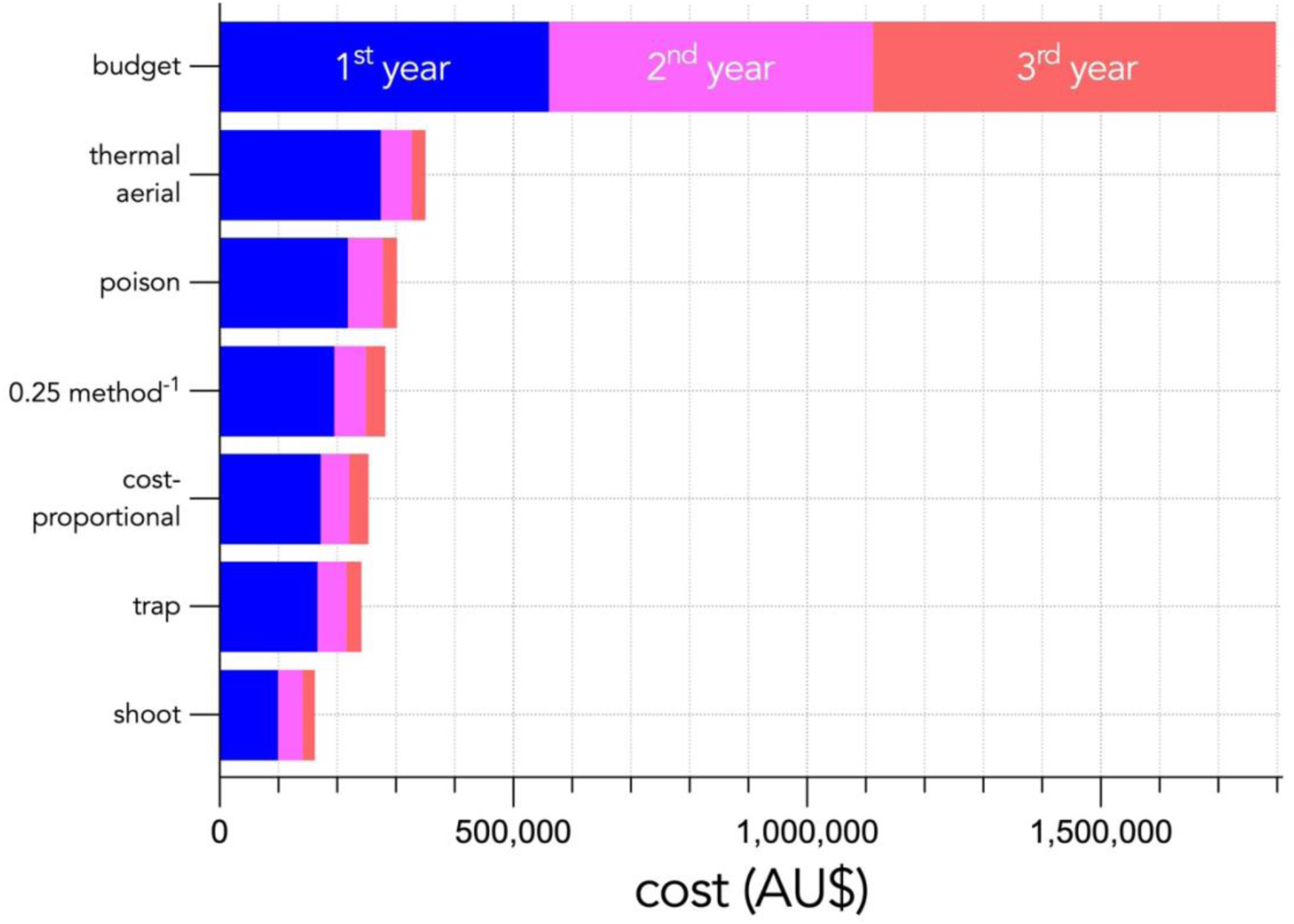
Median minimum cost per control scenario after 1, 2, and 3 years when applied at the most cost-effective annual harvest rate (0.95) to achieve reductions to ≤ 0.1*N*_0_. For comparison, ‘budget’ shows the cumulative operational budget allocated by the Government of South Australia over 1, 2, and 3 years to achieve feral pig eradication.

### Effort

For all harvest scenarios, minimum effort to reduce the population to ≤ 0.1*N*_0_ was achieved using an annual harvest of 0.95 (Fig. 5c,d). Thermal-assisted aerial culling required the lowest effort over 3- and 10-year projection intervals, with a minimum effort of 338 hours (291–396 hours) over 3 years (Fig. 5c), and 358 hours (298–436 hours) over 10 years (Fig. 5d).

Compared to thermal-assisted aerial culling, all other scenarios were relatively time-intensive. At the annual harvest with the lowest median total effort to achieve the reduction target over 3 years (0.95), estimated median effort for all other scenarios required between 644% (25% per harvest per method) and 870% (poisoning) more effort than thermal-assisted aerial culling (Fig. 5c,d). The 25% harvest per method and relative cost-proportional-allocation scenarios required the second- and third-lowest effort at all projection intervals above the minimum harvest rate (0.7) required to achieve the reduction target.

As with cumulative cost estimates above (Fig. 6), we derived estimates of median minimum effort over 1 and 2 years at the annual harvest of 0.95 to compare to median total effort after the 3-year projection interval (Fig. 7). After 1 year, median effort for thermal-assisted aerial culling was 282 hours (16% reduction) and 326 hours (3%) after two years compared to total median effort over the 3-year projection interval. We observed similar reductions for other cull scenarios, with reduction in effort ranging from 30% (shooting) to 16% (trapping) if culling stopped after 1 year, and 7% (shooting) to 3% (trapping) if culling stopped after 2 years.

**Figure 7.**
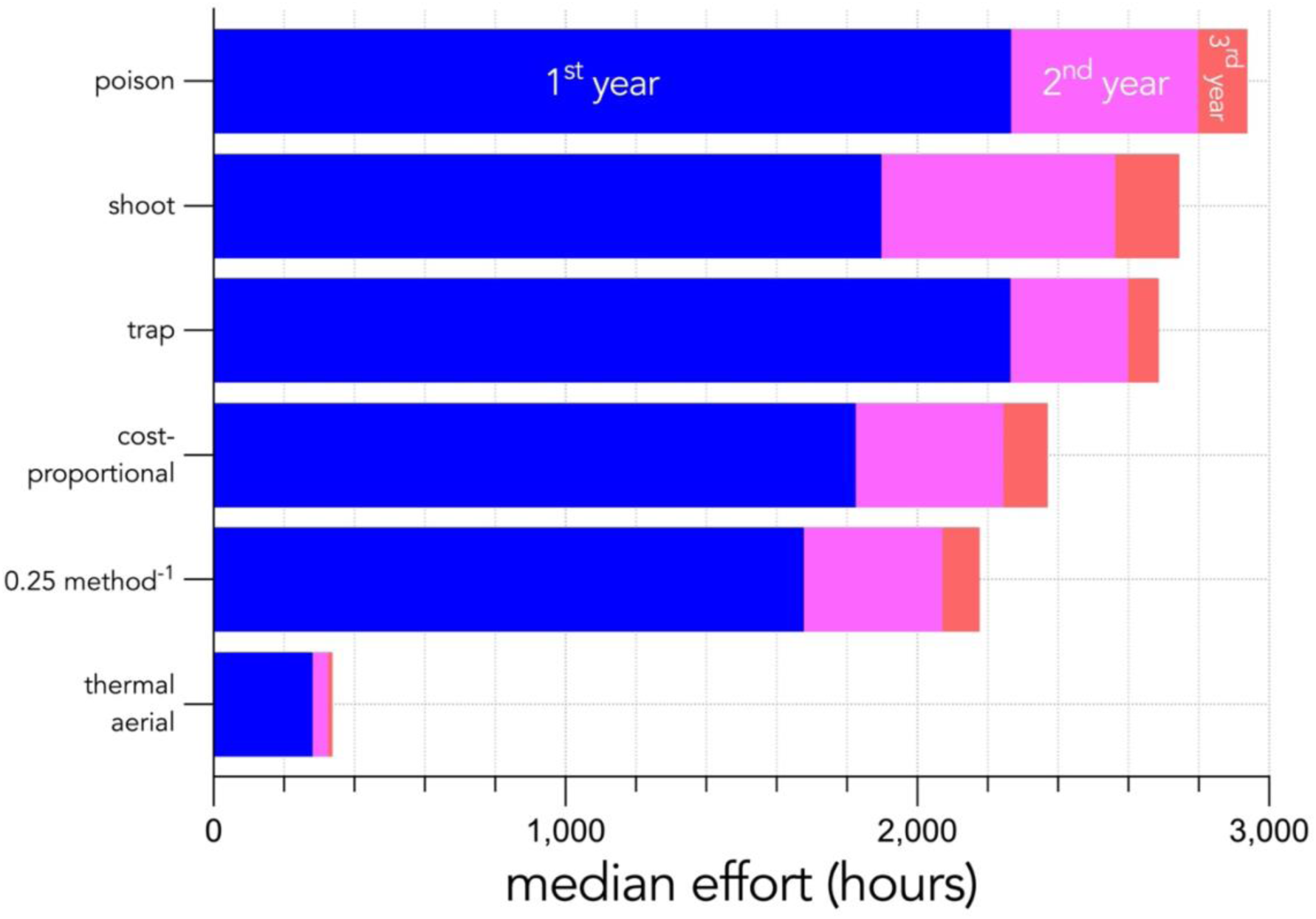
Median effort per year for each control scenario using the maximum annual harvest (0.95) over the 3-year projection interval.

## Discussion

Successful reduction of the pig population to ≤ 0.1*N*_0_ was achieved in all cull scenarios with annual harvest ≥ 0.7 over 3 years, or ≥ 0.5 over 10 years (Fig. 3). All simulations achieved the reduction target within the allocated budget (AU$2.66 million) at annual harvest ≥ 0.7 over 3 years (Fig. 5), which support the hypothesis that eradication is achievable within the project timeframe and budget.

We recommend a combination of all four control methods in accordance with current best practice (Massei et al. 2011), applied at the highest practical annual harvest in the first year. Effective distribution of methods will be dictated by landscape, pig density, and behaviour. While neither generic integrated pest-management scenarios achieved the lowest total median cost over either projection intervals (Fig. 5), both reduced population size to target sizes within the project budget and timeframe, as well as simulate realistic approaches compared to relying on a single culling method.

For all control scenarios, the maximum annual harvest was the most cost-effective strategy to achieve reduction targets (Fig. 5, 6). This outcome is logical because our models recalculated efficiency annually, so maximising offtake also maximises the number of pigs controlled at the highest efficiency (i.e., lowest effort pig^-1^). In reality, population density is not static throughout the year, but changes with every birth or death. Despite this simplification, our results agree with other studies reporting the benefits of rapid pig removal to achieve eradication based on the notion that rapid population reduction reduces potential for repopulation and development of avoidance behaviours (Cruz et al. 2005, McCann and Garcelon 2008, Parkes et al. 2010). Rapid control can also reduce the probability that other pressures might undermine success, such as waning public support, reduced staff motivation, and funding insecurity (Massei et al. 2011).

Thermal-assisted aerial culling required the lowest effort to achieve reduction targets compared to other control scenarios (Fig. 7), but it was the least cost-effective method (Fig. 5). The utility of thermal-assisted aerial culling appears greatest where rapid control is required, e.g., in the event of a serious livestock disease, or difficult-to-access areas where cost-effective control methods are impractical (see below). Thermal-assisted aerial culling is also useful in the final stages of pig eradication when the population is at low densities (Katahira et al. 1993), and cost is less important than the aim of eradication. Shooting was the least-expensive method because the model only accounts for labour and ammunition. In practice, shooting attracts additional costs including firearms, vehicles and fuel, labour and monitoring equipment. However, these additional costs are probably negligible compared to labour costs.

Programs that rely exclusively on ground-shooting are often unsuccessful (Keiter and Beasley 2017, Bengsen et al. 2020). An integrated pest-management approach using a combination of methods is considered the most effective because different strategies can be applied in response to changing densities, behaviours, and environmental conditions (Massei et al. 2011). For Kangaroo Island, large areas occupied by feral pigs cannot be accessed for shooting, trapping, or baiting, providing refuges from which reinvasion can occur (Hone et al. 1980, Choquenot 1996). Using thermal-assisted aerial culling in areas untreatable by other means appears justified, despite being the least cost-effective method for control. Incorporating assessment of habitat suitability and probability of pig occurrence (e.g., McMahon et al. 2010) might provide justification for these costs, but requires data describing patterns of food abundance and habitat use/movement. Without relevant local data, our aspatial projection is still useful for applying locally derived functional responses to compare cost and effort outcomes under different culling scenarios.

The realism of functional-response models could be improved with more accurate population estimates and reporting of operational effort. Systematic population estimates were unavailable, as were inferred changes in abundance. Efficiency data were not recorded for pig kills < 0.21 of the assumed total population (< 200 out of total 957) because the data-collection period ended when the estimated population fell below ∼ 200 pigs and the functional response was estimated based on the expected form in lieu of operational data. It is reasonable to assume that *per capita* effort required to remove pigs will continue to increase with decreasing pig density according to a Type II response, but the site-specific accuracy of the functional responses could be improved if data were available for lower population densities. Furthermore, only thermal-assisted aerial culling records included events with no pigs killed (i.e., *f_i_* > 0, *n_i_* = 0). Trapping and baiting effort was maintained until control outcomes were achieved and no unsuccessful trapping or baiting events occurred. However, it is possible that unsuccessful shooting events were unreported, increasing its apparent cost-effectiveness relative to other culling scenarios. Indeed, insufficient reporting on effort-outcome relationships (Hone et al. 2018) and imprecise accounting of program costs (Holmes et al. 2015) limit opportunity for analysis or improvement of efficiency.

Despite these limitations of data quality, using locally estimated functional responses is a valuable approach for advancing the accuracy of cost estimates compared to *ad hoc* estimation. For example, our modelled estimates predict that operational costs decrease substantially if high harvest is achieved in the first year (Fig. 6), contrasting with most funding schemes that allocate higher budgets in the final year of the program. The difference between actual allocated operational funding and estimated costs for the most cost-effective approach are in the order of > AU$1 million, representing a large savings that could be directed towards other environmental objectives.

### Conclusions

Applying locally estimated functional responses to estimate management costs at different population sizes and harvest rates is a novel approach that provides defensible insights into the relative cost-effectiveness of control methods. The real value of this approach is not in predicting the true costs, but rather in evaluating the relative costs of each scenario to identify the most cost-effective approach from a suite of available methods under different conditions stochastically within the bounds of the estimated model parameters. We were able to project realistic population change under varying harvest rates and produce realistic cost estimates to rank different control methods and scenarios currently used for pig eradication on Kangaroo Island, and to demonstrate the broader utility of this approach for informing decision-making in general pest animal management. Eradication of feral pigs on Kangaroo Island appears achievable within the project budget using sustained, high annual harvest, echoing approaches achieving successful pig eradication on islands elsewhere (e.g. Cruz et al. 2005, McCann and Garcelon 2008). Despite the potential for eradication on islands such as Kangaroo Island, the global feral pig distribution is predicted to expand (Lewis et al. 2017), and costs of biological invasions are projected to increase (Diagne et al. 2021). The use of operational control data to simulate management scenarios and maximise cost-effectiveness of control strategies will become increasingly necessary as pig distributions expand.

## Acknowledgements

We acknowledge the continued connection of Kaurna, Ramindjeri, and Ngarrindjeri people to Karta Pintingga (Kangaroo Island) and that this work was completed on the unceded lands of the Kaurna people. We pay our respects to Elders past, present and emerging. We thank the Biosecurity Division, Invasive Species Unit in the Department of Primary Industries and Regions South Australia for data access and support.

## Appendix S1: Comparing vital rates

Mortality rates for feral pigs in Australia ranges from 10–100% in juveniles and 15–50% in adults, while litter size ranges from 4.9–6.3 piglets, with sows producing 0.85 litters year^-1^ (Choquenot 1996). We calculated the mean and standard deviations of reported mortality ranges, assuming the range of juvenile mortality applied to age class *n*_1_ and adult mortality range applied to all other age classes (Table S1).

To calculate fertility, we divided the litter size range by 2 to give a range of females per litter, assuming a litter sex ratio of 1:1 (Snow et al. 2019). We then multiplied the female-only range by the average number of litters per year (*n* = 0.85) to produce a range of fertility from which we calculated the mean and standard deviation, as per age-specific survival probability. To allow for a small proportion of females to reproduce in their first year, we arbitrarily assumed a range of fertility for *n*_1_ that was 1/3 of the values used in all other age classes. This approach produced values comparable to those derived from Bieber and Ruf (2005). However, the frequency distribution within the reported ranges was unknown, so we considered the values derived for Australian feral pigs less reliable.

**Table S1:**
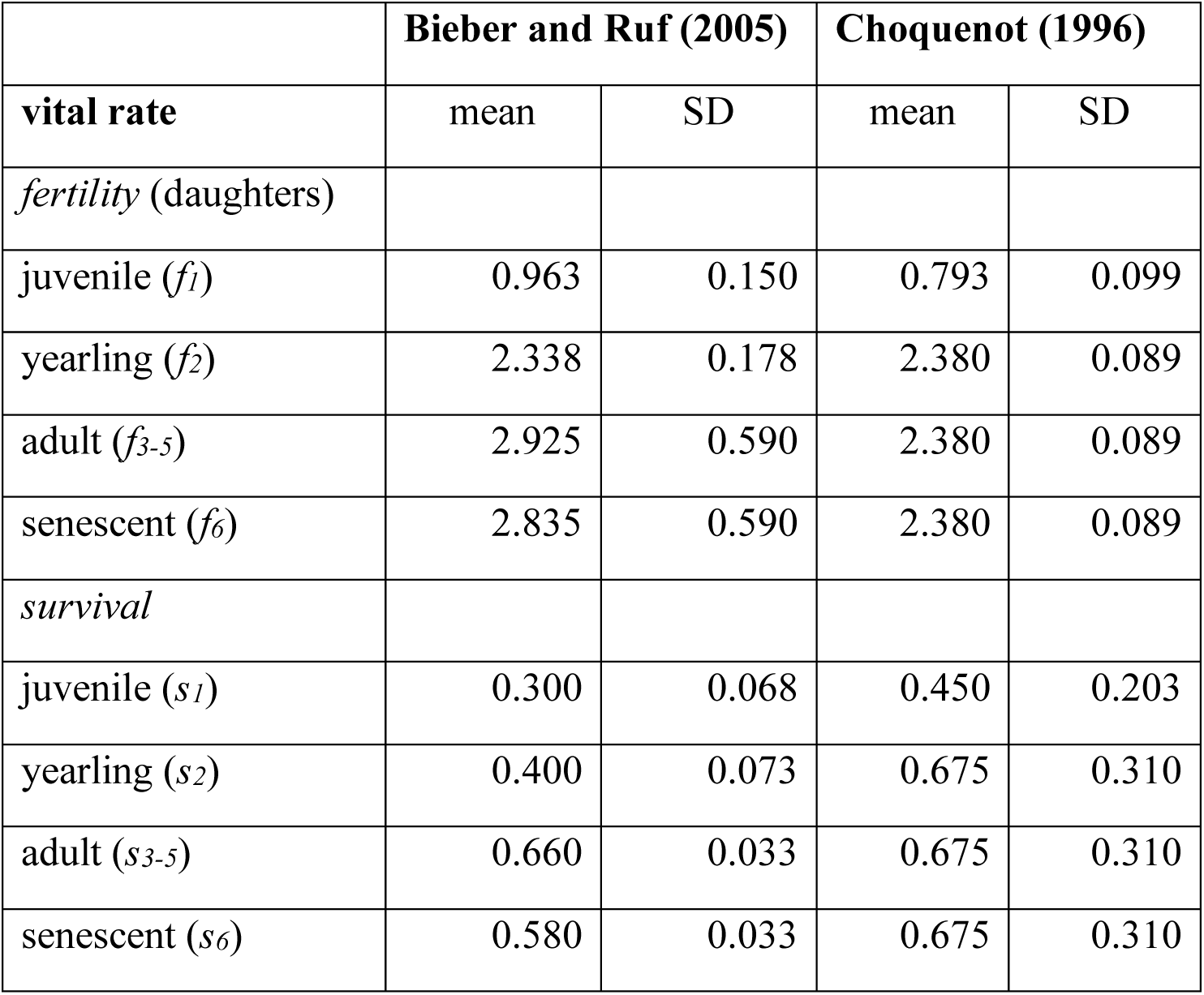
Comparison of age-specific vital rates (i.e., mean and associated standard deviation, SD) used in the model (Bieber and Ruf 2005) and those calculated from ranges reported for Australian feral pig populations (Choquenot 1996).

|  | Bieber and Ruf (2005) |  | Choquenot (1996) |  |
| --- | --- | --- | --- | --- |
| vital rate | mean | SD | mean | SD |
| *fertility* (daughters) |  |  |  |  |
| juvenile ($f_1$) | 0.963 | 0.150 | 0.793 | 0.099 |
| yearling ($f_2$) | 2.338 | 0.178 | 2.380 | 0.089 |
| adult ($f_{3-5}$) | 2.925 | 0.590 | 2.380 | 0.089 |
| senescent ($f_6$) | 2.835 | 0.590 | 2.380 | 0.089 |
| *survival* |  |  |  |  |
| juvenile ($s_1$) | 0.300 | 0.068 | 0.450 | 0.203 |
| yearling ($s_2$) | 0.400 | 0.073 | 0.675 | 0.310 |
| adult ($s_{3-5}$) | 0.660 | 0.033 | 0.675 | 0.310 |
| senescent ($s_6$) | 0.580 | 0.033 | 0.675 | 0.310 |

## Appendix S2: Stochastic population growth without Fmod

**Figure S1:**
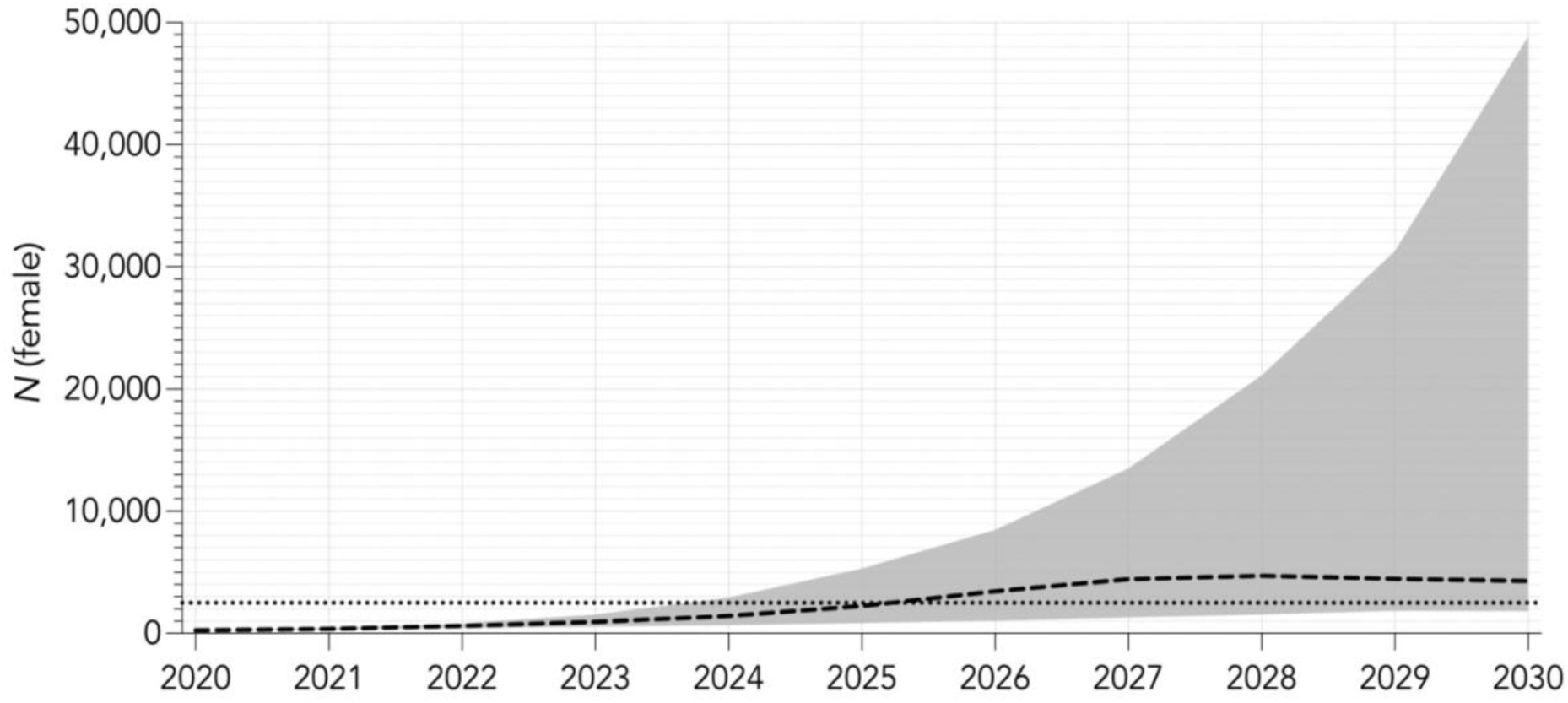
Stochastic population growth in an untreated population from an initial population (*N*_0_) of 250 females over 10-year projection interval with compensatory density feedback applied to survival only. Black dashed line indicates median population growth from 1000 iterations along with its upper and lower 95% confidence intervals (grey area), and black dotted line indicates estimated carrying capacity (*K*).

## Appendix S3: Control method-specific functional response coefficients

**Table S1:** Coefficients (α and β) for the exponential model fitted to each of the four control methods.

| control method | estimate |  |
| --- | --- | --- |
| | $\alpha$ | $\beta$ |
| shooting | 23.37 | 15.18 |
| thermal-assisted aerial culling | 1.45 | 6.43 |
| trapping | 10.97 | 5.83 |
| poison baiting | 18.04 | 10.87 |

## Appendix S4: Minimum annual cull rates and median estimated costs for reduction to < 0.01N0 in a population at carrying capacity over 3 and 10 years

We repeated the simulation in a population at carrying capacity (*N*_0_ = 2500) to compare minimum annual harvest rates and total median costs in the more likely management scenario where operational control begins when the pig population is at carrying capacity, as opposed to a population reduced by wildfire.

Using the same method described above, we determined that minimum annual harvests of 0.9 and 0.6 were required to achieve reductions to *N* ≤ 0.01*N*_0_ (≤ 25 females) over 3- and 10-year projection intervals (Fig. S2).

**Figure S2.**
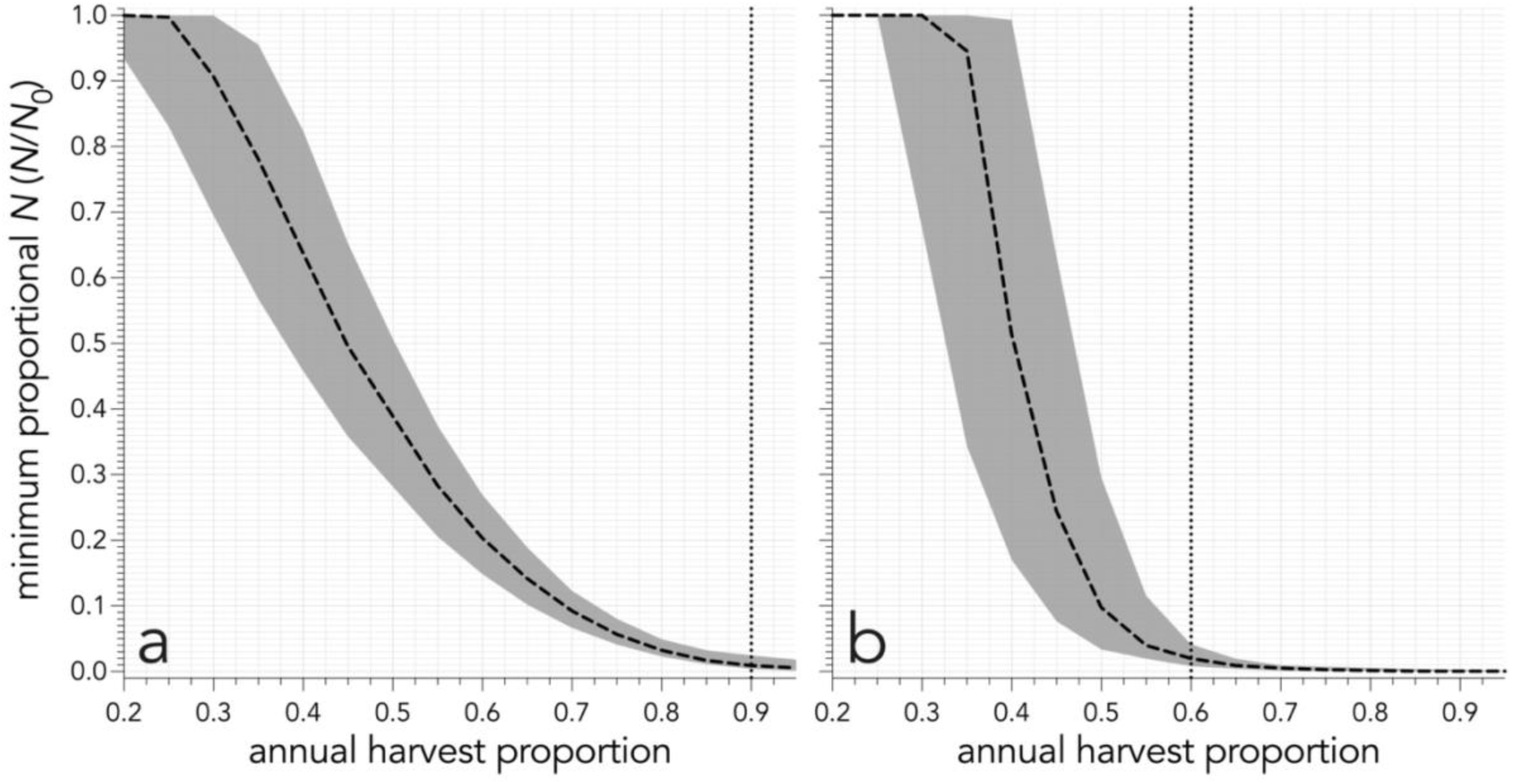
Proportion of initial population (*N =* 2500) remaining after (a) 3 years and (b) 10 years of annual harvest of 0.2 to 0.95, increasing in increments of 0.05. Harvest targets all age classes equally. Black dashed lines represent median proportion of population remaining along with its 95% confidence intervals (grey areas), vertical dotted lines represent harvest threshold required to reduce *N* to ≤ 0.01*N*_0_.

We refitted the exponential functional response model relative to the larger initial population size (*N*_0_ *= K*). We modified equation 4 to reflect changes in proportion of pigs remaining relative to *K*:

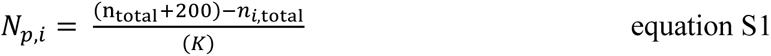

Similarly, we modified equation 6 such that the proportion of pigs remaining (*P*) was relative to *K*:

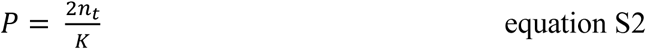

We repeated cull simulations for 1000 iterations over both 3- and 10- year projection intervals to produce estimates of total median cost (Fig. S3a,b) and effort (Fig. S4c,d) for each of the 6 cull scenarios. Over the 3-year projection interval, shooting had the lowest median cost when applied at an annual harvest rate of 0.9, whereas all other control scenarios produced their lowest total median cost when applied at the highest annual harvest of 0.95. Over the 10-year projection interval, all culling scenarios produced the lowest median total cost when applied at an annual harvest of 0.95.

When we applied an annual harvest of 0.95 to an initial population *N*_0_ *=* 2500, the reduction target was not met until the third year. Over the 3-year projection interval, cost savings could therefore not be achieved by suspending culling after only 1 or 2 years without compromising operational outcomes. However, substantial savings could be achieved relative to lowest total median costs over the 10-year projection interval when we applied culling scenarios at the annual harvest of 0.95, with operation suspended after 3 years when the reduction target (≤ 0.01*N*_0_) had been met. Using this approach, cost savings ranged from 20% (thermal-assisted aerial culling) to 42% (relative cost-proportional allocation) compared to the lowest total median costs if operation continued for 10 years.

**Figure S3.**
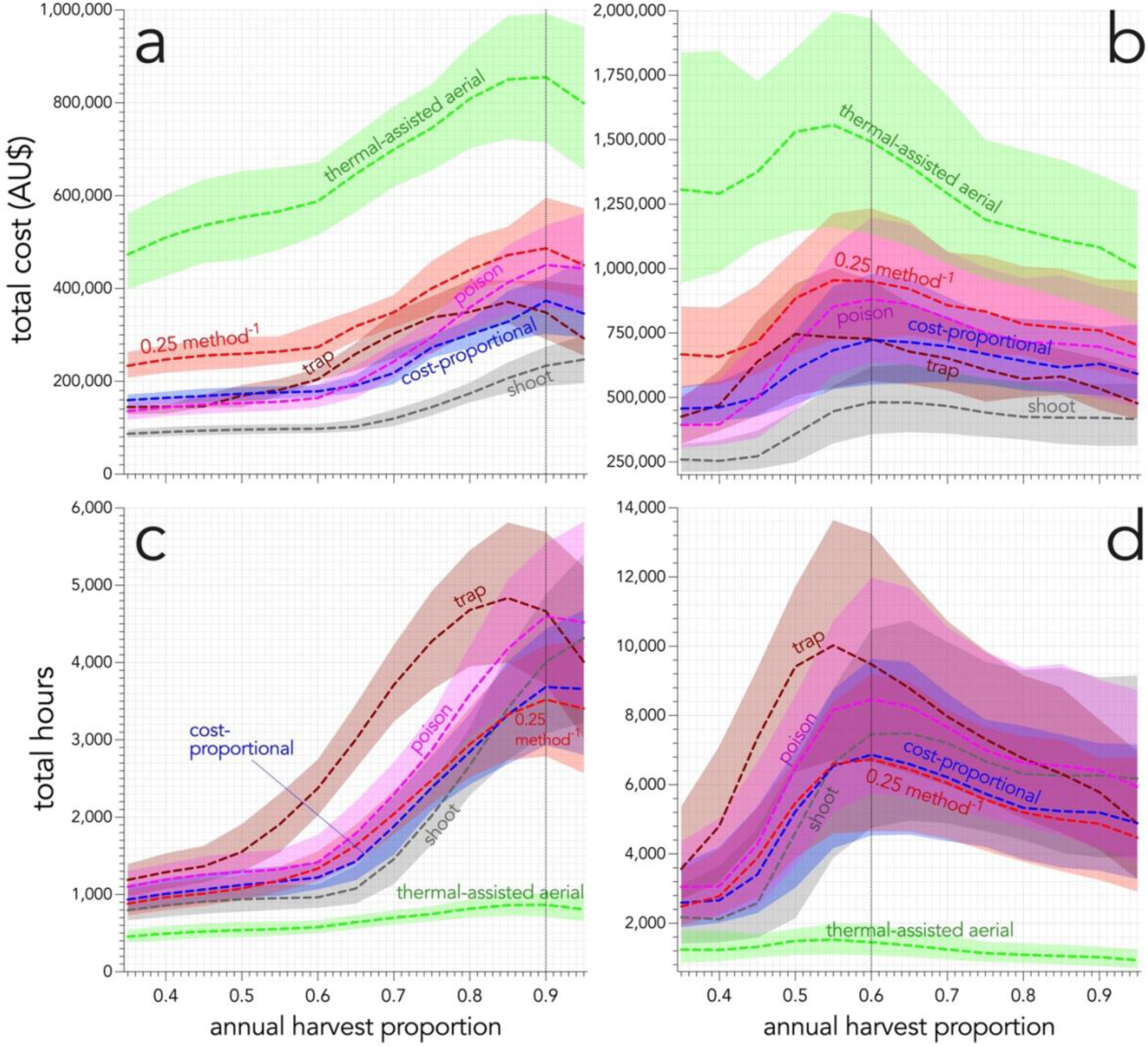
(a, b) Projected total cost (shaded areas = 95% confidence intervals) in a population *N*_0_ = 2500 over per cull scenario over (a) 3 and (b) 10 years, and (c, d) projected total effort (hours) per cull scenario over (c) 3 and (d) 10 years under increasing harvest proportions. Vertical dotted line represents minimum harvest required to achieve reduction to *N* ≤ 0.01*N*_0_. **thermal-assisted aerial** = thermal-assisted aerial culling; **0.25 method^-1^** = 25% harvest per method; **cost-proportional** = cost-proportional allocation.

Thermal-assisted aerial culling produced the minimum effort to reduce populations to the target size at annual harvest of 0.95 over both 3- and 10-year projection intervals (Fig. S3c,d). All other culling scenarios produced their lowest median total effort at an annual harvest of 0.95 over both 3- and 10-year projection intervals, except for shooting over 3 years that produced the lowest median total effort for that scenario and projection interval at an annual harvest of 0.9.

